# The evolutionary making of SARS-CoV-2

**DOI:** 10.1101/2021.01.29.428808

**Authors:** Ruben Iruegas, Julian Dosch, Mateusz Sikora, Gerhard Hummer, Roberto Covino, Ingo Ebersberger

**Affiliations:** Applied Bioinformatics Group, Inst of Cell Biology and Neuroscience, Goethe University Frankfurt, Frankfurt am Main, Germany; Department of Theoretical Biophysics, Max Planck Institute of Biophysics, Frankfurt am Main, Germany; Faculty of Physics, University of Vienna, Vienna, Austria; Institute of Biophysics, Goethe University Frankfurt, Frankfurt am Main, Germany; Frankfurt Institute for Advanced Studies (FIAS), Frankfurt am Main, Germany; Senckenberg Biodiversity and Climate Research Centre (S-BIK-F), Frankfurt am Main, Germany; LOEWE Centre for Translational Biodiversity Genomics (TBG), Frankfurt am Main, Germany

## Abstract

A mechanistic understanding of how SARS-CoV-2 (sarbecovirus, betacoronavirus) infects human cells is emerging, but the evolutionary trajectory that gave rise to this pathogen is poorly understood. Here we scan SARS-CoV-2 protein sequences in-silico for innovations along the evolutionary lineage starting with the last common ancestor of coronaviruses. SARS-CoV-2 substantially differs from viruses outside sarbecovirus both in its set of encoded proteins and in their domain architectures, indicating divergent functional demands. Within sarbecoviruses, sub-domain level profiling using predicted linear epitopes reveals how the primary interface between host cell and virus, the spike, was gradually reshaped. The only epitope that is private to SARS-CoV-2 overlaps with the furin cleavage site, a “switch” that modulates spike’s conformational landscape in response to host-cell interaction. This cleavage site has fundamental relevance for both immune evasion and cell infection, and the apparently ongoing evolutionary fine-tuning of its use by SARS-CoV-2 should be monitored.

## Introduction

Coronaviruses (CoV) are common human and animal pathogens (Narayanan, et al. 2015; Weiss and Navas-Martin 2005), which had historically been associated with only considerably mild symptoms. Following two global CoV outbreaks associated with severe acute respiratory syndromes (SARS) caused by SARS-CoV-1 (2002) and MERS-CoV (2012), respectively, CoVs are acknowledged as potent emerging human pathogens (Coleman and Frieman 2014). With SARS-CoV-2 as the causative agent of COVID19 (Zhou, et al. 2020b), the first coronavirus has entered the top ranks of human viral pathogens. In only 12 months, the WHO reported worldwide 85.5 million confirmed infections and 1.8 million deaths (https://covid19.who.int), and the socio-economic impact is unprecedented.

SARS-CoV-2 and SARS-CoV-1 belong to the subgenus sarbecovirus within the beta-coronaviruses (Coronaviridae Study Group of the International Committee on Taxonomy of 2020). Sarbecoviruses are prevalent in bats (Hu, et al. 2017; Li, et al. 2005), and it is general consensus that horseshoe bats act as natural reservoirs of these viruses (Cui, et al. 2019; Zhou, et al. 2020b). While SARS-CoV-1 presumably used civets as a jumping host (Guan, et al. 2003; Li, et al. 2005; Wu, et al. 2005), the closest known relatives to SARS-CoV-2 are two bat viruses. RaTG13 was isolated already in 2013 from the intermediate horseshoe bat, *Rhinolophus affinis* (Zhou, et al. 2020b), and only recently RmYN02 was identified via a metagenomics sequencing of viruses isolated from the Malayan horseshoe bat, *R. malayanus* (Zhou, et al. 2020a).

The SARS-CoV-2 genome comprises 10 non-overlapping open reading frames (ORFs; Figure 1A). Four ORFs encode the structural building blocks of the virus, the nucleocapsid phosphoprotein (N), the membrane glycoprotein (M), the surface glycoprotein (spike), and the envelope protein (E). A fifth ORF holds the information for the polyproteins 1a and 1ab, which are post-translationally cleaved into 16 sub-units comprising the replication machinery of the virus (de Wit, et al. 2016; Narayanan, et al. 2015; Ziebuhr, et al. 2000). The precise functions of the proteins encoded by the remaining ORFs are only partly understood, and it is assumed that they play a role in fine-tuning host interaction (Issa, et al. 2020; Li, et al. 2020a; Liu, et al. 2014; Miorin, et al. 2020; Morante, et al. 2020).

**Figure 1.**
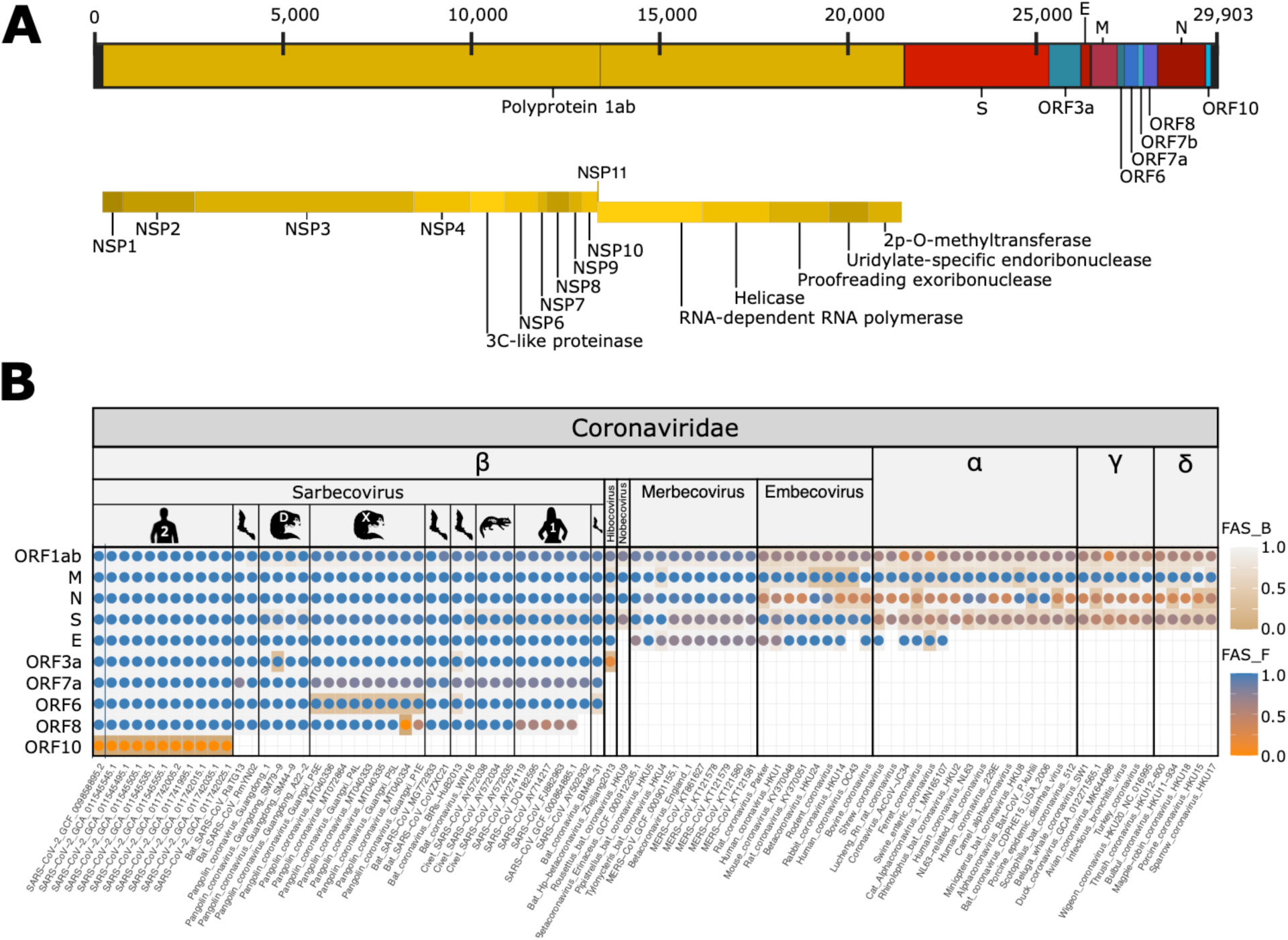
The phylogenetic profile of the SARS-CoV-2 protein set. **(A)** Position of open reading frames (ORFs) in the genome of SARS-CoV-2. Represented are the polyprotein and its components (yellow), structural proteins (red), and accessory proteins (blue). **(B)** Feature-aware phylogenetic profiles of SARS-CoV-2 proteins across the Coronaviridae. Sarbecoviral hosts are indicated by the pictograms. Human 1 and 2 represent SARS-CoV-1 and SARS-CoV-2, respectively. The Guangdong and Guangxi pangolin viruses are marked with ‘D’ and ‘X’, respectively. Taxa are ordered with increasing evolutionary distance to SARS-CoV-2 GCF_009858895.2, which was used as the seed species in this analysis. Dots indicate the detection of an ortholog in the respective species. Colors represent the feature architecture similarity between the seed protein and its ortholog using once the seed (dot color; FAS_F), and once the ortholog (cell color; FAS_B) as reference. While feature architectures are largely conserved within sarbecoviruses, we see pronounced differences indicated by low FAS scores in merbecoviruses, embecoviruses, and outside the beta-CoV. Abbreviations: M – membrane glycoprotein, N – nucleocapsid protein, S – spike glycoprotein, E – envelope protein.

Of all proteins encoded in the SARS-CoV-2 genome, the spike has received most attention. The 1273 amino acid long, heavily glycosylated protein (Walls, et al. 2020; Walls, et al. 2016) mediates the primary interaction of the virus with its host (Walls, et al. 2020; Walls, et al. 2017) and thus is the main target of the human immune response (Shrock, et al. 2020). Like its ortholog in SARS-CoV-1 (Lan, et al. 2020; Li, et al. 2003) the spike interacts via its receptor binding domain (RBD), and the receptor binding motif (RBM) therein, with the angiotensin-converting enzyme 2 (ACE-2) receptor of the host cell (Lan, et al. 2020; Letko, et al. 2020; Walls, et al. 2020; Wrobel, et al. 2020). The spike is organized as a homo-trimer (Turonova, et al. 2020; Wrapp, et al. 2020). In each chain, the RBD can either point down or up. Spike is in a ‘closed’ state when all RBDs are down, and it is ‘open’ when at least one RBD is up (Wrobel, et al. 2020). During the infection process, spike can be proteolytically cleaved into the receptor binding domain (S1) and the fusion domain (S2) (Hoffmann, et al. 2020b; Jaimes, et al. 2020; Shang, et al. 2020a) by furin, a human endoprotease active at multiple compartments between the trans-Golgi network and the cell surface (Braun and Sauter 2019; Molloy, et al. 1999). The resulting destabilization of the trimer shifts the balance of the receptor conformation towards an open state, which increases the binding affinity to human ACE2 by three orders of magnitude (Wrobel, et al. 2020). Proteolytic cleavage by the TMPRSS2 protease activates the fusion of the viral envelope with host membranes (Hoffmann, et al. 2020a; Hoffmann, et al. 2020b; Shang, et al. 2020a; Walls, et al. 2017). Although the initial furin cleavage is not strictly required for the infection process (e.g. (Turonova, et al. 2020)), it enhances cell entry (Shang, et al. 2020a). Structural analyses of the spike and of its interaction with ACE2 have meanwhile revealed the position and identity of the residues in the receptor-binding domain that are responsible for the tight binding between the two proteins (Lan, et al. 2020; Liu, et al. 2020; Mehdipour and Hummer 2020; Shang, et al. 2020b; Wrobel, et al. 2020).

Among all known animal sarbecoviruses, the RBM of the pangolin virus spike shows the highest similarity (98.6%) to RBM of SARS-CoV-2 (Liu, et al. 2020). Since pangolins are known reservoirs for beta-coronaviruses, it was suspected that a recombinant between a pangolin virus and a bat virus gave rise to SARS-CoV-2 (Lam, et al. 2020; Li, et al. 2020b; Liu, et al. 2020; Xiao, et al. 2020). However, more recently evidence accumulated that RaTG13 secondarily modified the RBM in its spike (Boni, et al. 2020). This does not preclude pangolins as the jumping host, but evidence in favor of a direct transmission from bats has accumulated (Boni, et al. 2020).

Since the first reported case of Covid-19 in late 2019, the understanding of how SARS-CoV-2 interacts with the human host, and how the virus evolves in the human lineage (Korber, et al. 2020) has substantially increased. Yet, the evolutionary trajectory that turned an animal virus into a potent human pathogen remains obscure. Here we present a high-resolution phylogenetic profiling analysis of the proteins encoded in the SARS-CoV-2 genome across representatives covering the diversity of the coronaviridae. Our results reveal that sarbecoviruses are distinct from other members of the beta-coronaviruses both with respect to the set of encoded proteins and of their domain architectures. On this level of resolution, changes within the sarbecoviruses are scarce and provide no information about the emergence of SARS-CoV-2. This picture changes when we use predicted linear T-cell epitopes to scan the spike surface for genetic innovation. Here, we see a prominent signal pointing towards a gradual reshaping of spike during virus evolution, which sheds light on the evolutionary making of this pathogen.

## Methods

### Data collection

A collection of 279 virus genomes covering the currently known diversity of coronaviridae was downloaded from the RefSeq, GenBank, ViPR, and GISAID databases (Shu and McCauley 2017). The taxon sampling includes four subgenera of the beta-coronaviruses, i.e., sarbecovirus harboring SARS-CoV-2 and SARS-CoV-1, together with closely related strains isolated from pangolins, bats and civets, hibecovirus, nobecovirus, merbecovirus, and embecovirus. Representatives from the alpha-, gamma-, and delta-coronaviridae complement the taxon sampling. The list of taxa is provided in Table S1.

### Phylogenetic profiling and feature architecture comparison

Phylogenetic profiles of the 10 ORFs annotated in the genome of SARS-CoV-2 were generated with the targeted ortholog search tool fDOG (www.github.com/BIONF/fDOG) (Ebersberger, et al. 2014) using the strain ASM985889v3 (RefSeq Assembly accession GCF_009858895.2) as the seed. Linear protein features were annotated in the following way: Pfam (Pfam v.33.1) (El-Gebali, et al. 2019) and SMART (Letunic, et al. 2009) domains were annotated with hmmsearch from the HMMER package (Finn, et al. 2015), low complexity regions were predicted with flps (Harrison 2017) and SEG (Wootton and Federhen 1993), signal peptides with SignalP (Petersen, et al. 2011), transmembrane domains with tmhmm (Sonnhammer, et al. 1998), and coiled-coil conformations with COILS2 (Lupas, et al. 1991). The resulting feature architectures (FA) were compared pairwise between the seed proteins and each of their orthologs, and feature architecture similarity scores (FAS scores)(Koestler, et al. 2010) were computed with GreedyFAS v.1.4.8 (https://github.com/BIONF/FAS) integrated into the fDOG package. FAS score ranges between 0 (FA_seed_and FA_ortholog_are disjunct) and 1 (FA_seed_= FA_ortholog_). Feature-aware phylogenetic profiles were visualized and analyzed with PhyloProfile v.1.4 (Tran, et al. 2018).

### Phylogenetic tree reconstruction

For maximum likelihood (ML) tree reconstruction, multiple sequence alignments of the orthologous groups for S, N, and ORF1ab were created with mafft-linsi (Katoh and Toh 2008), respectively. ML trees for the individual alignments were computed with RAxML (Stamatakis 2014), allowing the software to automatically select the best fitting substitution model (Option PROTGAMMAAUTO). Branch support was assessed with 100 bootstrap replicates. The viral phylogeny was then computed from the individual gene trees with ASTRAL III (Zhang, et al. 2018).

### Epitope prediction

Spike proteins were scanned for the presence of linear T-cell epitopes using the consensus epitope prediction approach (Moutaftsi, et al. 2006; Wang, et al. 2008) provided via the Immune Epitope Database website (http://www.iedb.org/)(Vita, et al. 2019). Major histocompatibility complex (MHC) anchor residues in the antigen were predicted by using the models based on reference sets for class I (Greenbaum, et al. 2011) and II (Weiskopf, et al. 2013). The two sets were compiled such that the represented alleles cover > 97% (MHC-I) and >99% (MHC-II) of the general population provided by the Allele frequency net database (Gonzalez-Galarza, et al. 2020). Linear epitopes were filtered using a percentile rank cutoff of 0.1 and considered homologous if the aligned region in the orthologous sequence predicts binding affinity to the same MHC allele. Epitopes were classified according to their node of innovation and visualized on a 3D model of the full length, glycosylated SARS-CoV-2 spike protein embedded in a membrane (Sikora, et al. 2020) (publicly available at 10.5281/zenodo.4442942), using the Visual Molecular Dynamics (VMD) software v.1.9 (Humphrey, et al. 1996).

### Surface accessibility analysis

Surface accessibility of the spike was reported in (Sikora, et al. 2020), and is based on an extensive atomistic molecular dynamics (MD) simulation of a viral envelope patch containing 4 spike proteins in realistic conditions. The analysis considers the spike’s structural flexibility and the steric dynamic shielding by the glycans that cover the spike surface. Accessibility analyses were done once with spikes in the conformation with 2 RBDs down and one up, and once with all RBDs down. The analysis’ raw data is publicly available on 10.5281/zenodo.4442942.

## Results

We used an evolutionary approach to shed light on the trajectory that ultimately gave rise to SARS-CoV-2. The phylogenetic profiles of the SARS-CoV-2 proteins across coronaviridae (Fig. 1) reveal a coronavirus core genome comprising only four ORFs (M, N, S, and the P1). We find orthologs to the envelope protein, an essential structural and functional constituent of the shell of all coronaviruses (Mukherjee, et al. 2020; Schoeman and Fielding 2019), only in the alpha- and beta-CoV. However, identifying distant orthologs for short proteins—the envelope protein has a length of only 75aa—is difficult (Jain, et al. 2018), and even more so since a conserved domain that is characteristic for the envelope protein in alpha- and beta-CoV is missing in the envelope protein of gamma- and delta-CoV (Schoeman and Fielding 2019). Thus, a limited sensitivity of the ortholog search (Jain, et al. 2019) likely explains why we failed to establish orthology relationships between the envelope proteins across all coronaviridae. A multi-gene phylogeny (Fig. 2) recovers the four genera alpha-, beta-, gamma- and delta-CoV as monophyletic clades, respectively, and midpoint rooting places the alpha- and beta-coronaviruses as sister clades. Within beta-CoV, embecoviruses branch off first, followed by the merbecoviruses, which harbors with MERS the so far most aggressive human pathogen within the beta-coronaviruses (Petersen, et al. 2020). Nobecovirus and hibecovirus represent the closest relatives of sarbecovirus. Sarbecoviruses themselves are represented by three distinct lineages. The bat coronavirus BM48-31 branched off prior to the diversification of the monophyletic clades harboring SARS and SARS-CoV-2, respectively. Within the latter clade, the bat virus RaTG13 is placed stably as the closest known relative to SARS-CoV-2, followed by RmYN02, and the paraphyletic pangolin viruses from Guangdong and Guangxi, respectively. Two further bat viruses represent the early branching lineages in this clade. Considering the distribution of viral hosts, the phylogeny reveals that only bat viruses cover the full phylogenetic diversity of sarbecoviruses (Fig. 2).

**Figure 2.**
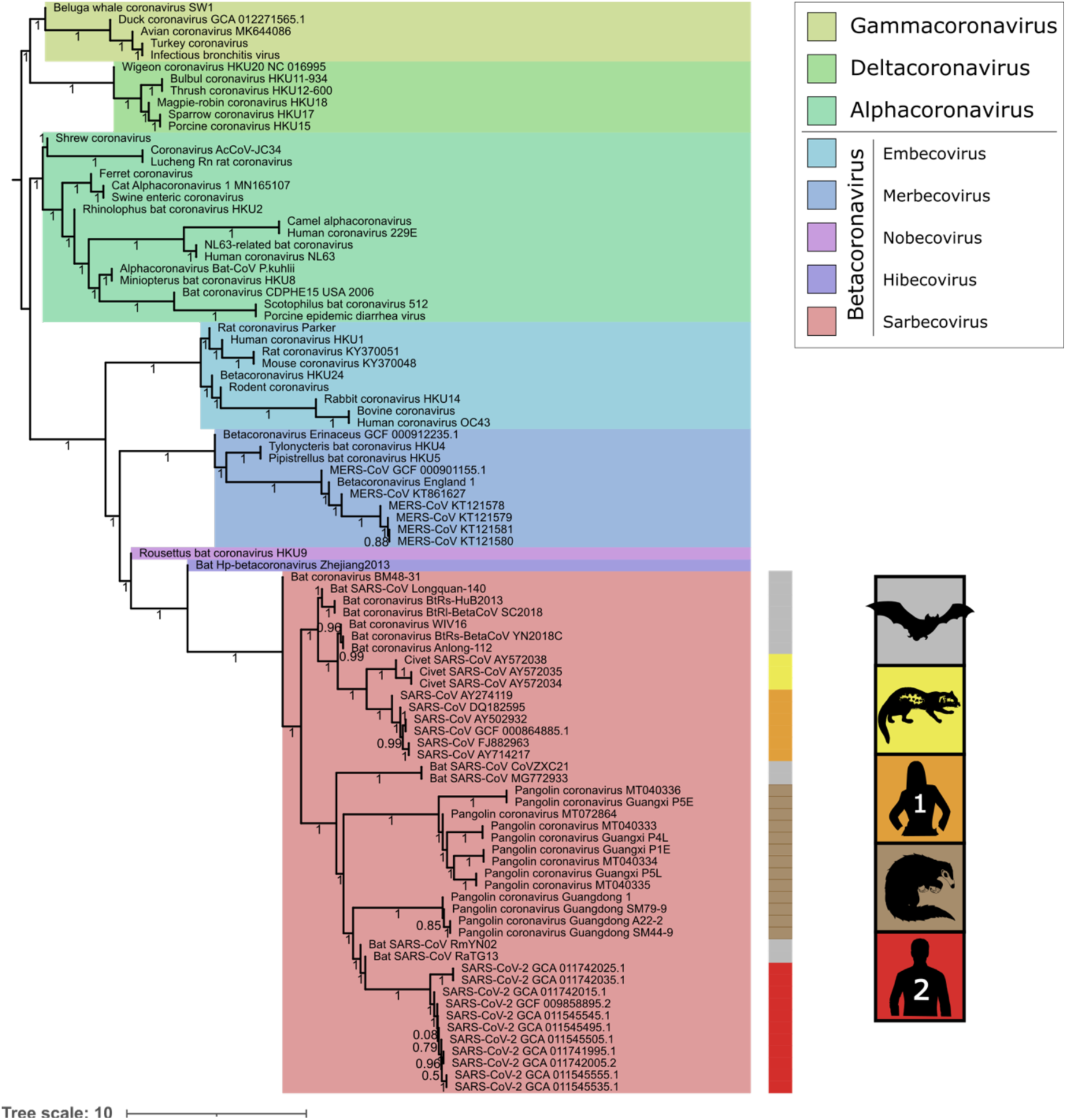
Phylogenetic relationships within the coronaviridae. Colored areas in the tree indicate the different genera within coronaviridae. The beta-coronaviruses are resolved to their respective sub-genera. The colored bar next to the sarbecovirus clades denotes the host species from which the viruses were sampled. Clade-specific hosts within sarbecovirus are indicated by pictograms, where the human viruses 1 (orange) and 2 (red) represent SARS-CoV-1 and SARS-CoV-2, respectively. The tree supports the genera and sub-genera nested within CoV as monophyletic clades and supports hibecovirus and nobecovirus as the closest relatives of the sarbecoviruses, the sub-genus comprising both SARS-CoV-1 and SARS-CoV-2.

Arranging the presence/absence pattern of orthologs to the SARS-CoV-2 proteins according to the reconstructed tree shows that sarbecoviruses are not only phylogenetically but also genetically distinct from other coronaviruses (cf. Fig. 1B). ORFs 6 - 10 are confined to this sub-genus. Only ORF3a, which was previously considered as unique to sarbecovirus (Issa, et al. 2020) has an ortholog at the correct position between the spike and the envelope protein also in the hibecovirus representative. The emergence of the sarbecovirus sub-genus during beta-coronavirus diversification coincides with a complete exchange of the accessory gene set. Within sarbecovirus the gene set is stable. ORF10 encoding a 38 amino acid long polypeptide, whose orthologs are confined to SARS-CoV-2 seems a notable exception. Yet, a closer look revealed that almost identical sequences exist in representatives of all sarbecovirus lineages (100% coverage, 99% - 82% id; Fig. S1), which are not annotated as an ORF. It was proposed that ORF10 might be a spurious gene prediction because no transcript could be detected in RNAseq data (Kim, et al. 2020).

However, in the light of its maintenance and conservation throughout sarbecoviral evolution this assessment should be re-considered. In either case, the alleged gene gain of SARS-CoV-2 reduces to a difference in gene annotation.

### Evolution of protein feature architecture

The remarkably stable gene set reveals that neither the acquisition nor the loss of genes is a driving force of sarbecovirus evolution. We therefore increased the resolution to the level of linear features annotated along the individual proteins, such as Pfam domains, transmembrane regions, secondary structures or low complexity regions. The comparison of the resulting feature architectures (Koestler, et al. 2010) between orthologs revealed a more differentiated evolutionary pattern. Among the beta-CoV core proteins, we see lineage-specific variations in their architectures on the genus and sub-genus level for three proteins (E, N, and S). The differences are mainly due to the differential presence of regions enriched with a particular amino acid, and of secondary-structure elements (see Fig. S2 for the spike). In particular, terminal low complexity regions are often involved in protein-protein interactions (Coletta, et al. 2010), and it will be interesting to see whether the differences detected here do have functional consequences. For the non-structural proteins (NSPs) contained in the polyprotein 1ab, we largely see a similar picture (Figs. S3, Table S2), where overall differences are mainly due to lineage-specific gains or losses of low complexity regions. This agrees with the notion that the enzymatic activity and functional domains of essential NSPs are conserved across CoV diversity (Snijder, et al. 2003; Thiel, et al. 2003). However, there are two notable exceptions, NSP1 and NSP3.

NSP1 is involved in shutting down the translation of host mRNAs to free capacities for the production of viral proteins (Narayanan, et al. 2015; Tanaka, et al. 2012). The phylogenetic profiling yielded NSP1 orthologs only in beta-CoV but not in the other genera. This is in line with previous findings that alpha- and beta-CoV NSP1s share no significant sequence similarity (Connor and Roper 2007; Jansson 2013), and with the absence of NSP1 in gamma- and delta-CoV (Snijder, et al. 2003; Woo, et al. 2010). Within beta-CoV, no orthology relationships could be established to MERS NSP1. Despite the existence of individual conserved motifs (Fig. S4), MERS NSP1 has diverged beyond recognition as an ortholog. This pronounced change in sequence goes hand in hand with a different mode of action. Other than NSP1 of SARS-CoV-1, MERS NSP1 does not form a stable complex with the 40S ribosomal subunit to block the translation of host mRNAs. Instead, it inhibits selectively the translation of mRNAs transcribed in the nucleus but not of viral mRNAs that are transcribed in the cytoplasm (Lokugamage, et al. 2015). NSP3, the second protein with a notable change in its feature architecture, is a large transmembrane protein. It forms the core of a pore complex that spans both membranes of the cytosolic vesicle where viral genome replication takes place (Wolff, et al. 2020). NSP3 is suspected to play a key role in the transfer of messenger RNAs from the replication compartments to the cytosol. A comparison of the feature architectures for this protein reveals two major changes during beta-CoV evolution (Fig. S5). NSP3 of embecovirus, as well as of all alpha-, gamma-, and delta-CoVs in our analysis harbor two CoV_peptidase domains representing papain-like protease (PL^Pro^) domains that are used to release NSP3 from the polyprotein (Lei, et al. 2018). The N-terminal peptidase domain, PL^Pro^-1 was lost after the split of the embecovirus lineage. This finding is in line with previous reports (Lei, et al. 2018; Neuman 2016), but the functional implication of this domain loss is still unclear. The second major change is the gain of the Pfam domain bCoV_SUD_C (PF12124) in the last common ancestor of the sarbecoviruses. This domain is involved in the binding of single-stranded RNA (Johnson, et al. 2010). Its emergence in the sarbecoviruses provides an initial indication that the precise ways how mRNA export is mediated has been changed during beta-coronavirus evolution. Orthologs of the SARS-CoV-2 accessory proteins are, with one exception, absent outside sarbecovirus and their evolutionary origins remain elusive. The feature architecture comparisons of ORF3a, which is represented by an ortholog in hibecovirus, provides unprecedented insights into the evolutionary origins of this protein (Fig. 3). ORF3a forms a homo-tetrameric ion channel (Issa, et al. 2020; Lu, et al. 2006), it is pro-apoptotic and localizes at the endoplasmic reticulum – Golgi compartment (Issa, et al. 2020; Minakshi, et al. 2009). The sarbecovirus orthologs consistently display 3 transmembrane domains in their N-terminal half, together with a bCoV_viroporin Pfam domain (PF11289) spanning the full protein. The hibecovirus ortholog of ORF3a displays also three transmembrane domains. Instead of the bCoV_viroporin domain, its central part displays a weak similarity to the CoV_M Pfam domain (PF01635; e-value: 0.25). This domain is otherwise characteristic for the membrane protein M, which interacts with the spike and the envelope protein to form the virus envelope at the endoplasmatic reticulum – Golgi intermediate compartment (ERGIC, (Ujike and Taguchi 2015)). Notably, the length of M is similar to that of ORF3a in sarbecovirus and hibecovirus, it shares with ORF3a of both sub-genera the presence of three transmembrane domains in its N-terminal half, and with hibecovirus ORF3a the presence of the CoV_M Pfam domain. Eventually, both M and ORF3a act at the same sub-cellular localization, the ERGIC. Together, this lets us hypothesize that ORF3a emerged by a duplication of M in the last common ancestor of sarbecovirus and hibecovirus and subsequently diversified in sequence, domain architecture, and function.

**Figure 3.**
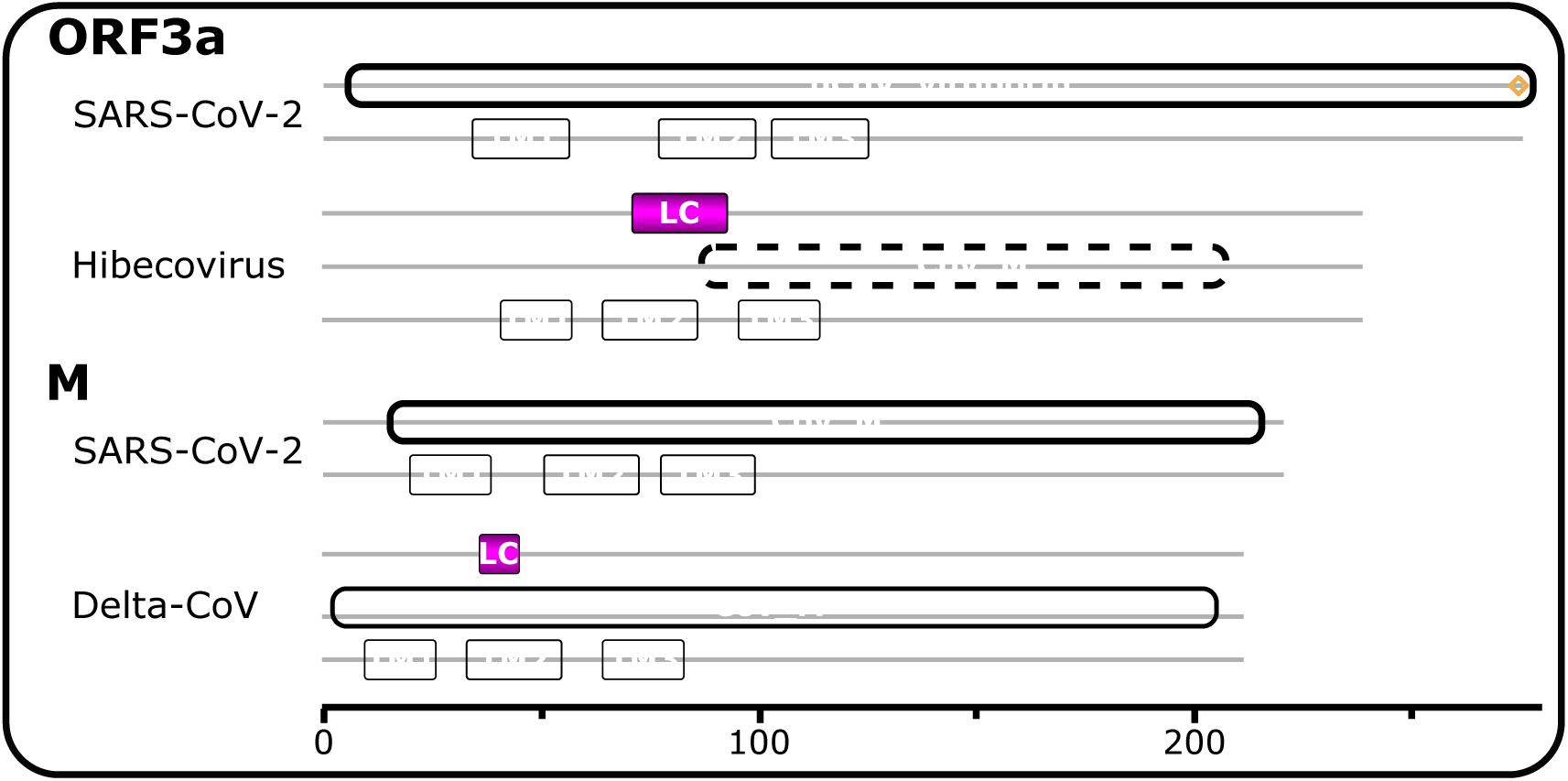
Feature architectures of SARS-CoV-2 ORF3a and M, and of their evolutionarily most distantly related CoV orthologs. Grey lines represent the amino acid sequences of the respective proteins, and colored bars indicate position and length of the protein parts occupied by the corresponding features. bCoV_viroporin and CoV_M represent the Pfam domains PF01635 and PF11289, respectively. The CoV_M domain in hibecovirus ORF3a (hatched box) was identified with an e-value of 0.24. TM1-3 represent three transmembrane helices, and LC the position of regions with a biased amino acid composition. ORF3a SARS-CoV-2: YP_009724391.1; ORF3A Hibecovirus: YP_009072441.1; M SARS-CoV-2: YP_009724393.1; M Delta-CoV: YP_005352848.1

Similar to the findings on the gene level, we observe for the entire gene set remarkably conserved feature architectures within the sarbecoviruses (cf. Figs. 1B and S3). Of the three lineage-specific changes, two represent losses of low-complexity regions in the proteins encoded by ORFs 6 and 7A. The third event, however, is a loss of an N-terminal signal peptide in SARS-CoV-1 due to a 39 amino acid long deletion (Fig. S6) that is otherwise prevalent in the sarbecoviruses. The protein encoded by ORF8 is involved in envelope protein regulation, cell death, viral replication, and suppression of the interferon response from the host (Wong, et al. 2018). ORF8 deletions in SARS-CoV-1 have been associated with attenuated replication, probably as a consequence of the virus’ adaptation to the human host (Muth, et al. 2018). Meanwhile, lineage-specific deletions in ORF8 emerged also in SARS-CoV-2 (Gamage, et al. 2020; Rambaut, et al. 2020; Su, et al. 2020), which suggests that ORF8 might be a general way to fine-tune virus activity within the human host.

### Evolution of the epitope landscape

The comparison of feature architectures between SARS-CoV-2 proteins and their orthologs within and outside sarbecorvirus revealed a number of differences, likely with functional consequences. However, there is still no indication that any of the underlying changes have contributed to the evolutionary emergence of SARS-CoV-2. Spike, while modified in the course of sarbecovirus evolution (Jaimes, et al. 2020; Lan, et al. 2020; Walls, et al. 2020; Wrobel, et al. 2020; Zhou, et al. 2020a), displays a fully conserved domain architecture within sarbecovirus. We therefore scanned the spike for changes on the sub-domain level. Epitopes are short structural elements exposed on the protein surface that are visible to, and recognized by, antibodies and other proteins of the host. The epitope landscape of the spike should therefore reveal the interface via which the virus interacts with the host, not only in eliciting immune response (Ng, et al. 2020; Shrock, et al. 2020; Zhang, et al. 2020a) but also during the infection process.

The consensus epitope prediction approach (Moutaftsi, et al. 2006; Wang, et al. 2008) resulted in nine linear epitopes (LE) along the sequence of the SARS-CoV-2 spike (Figs. 4A and S7). With the exception of LE3, all are located in the S1 domain of the spike, and LE9 identifies the furin cleavage site that separates S1 from S2. LE2,4-6 reside in the RBD, and LE6 overlaps with the RBM therein.

**Figure 4.**
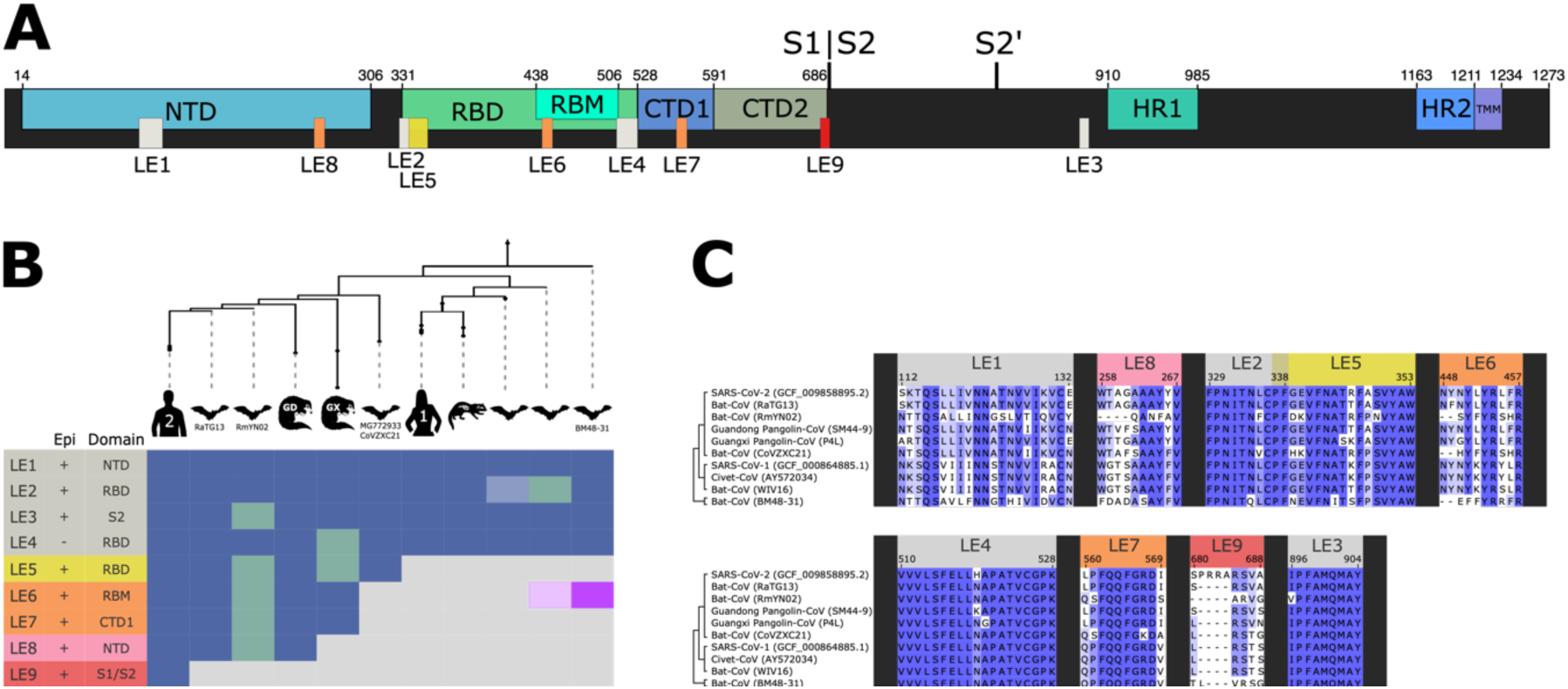
The phylogenetic profiles of linear epitopes in the spike glycoprotein of SARS-CoV-2. **(A)** Position of domains and linear epitopes in the spike. Represented are the N-terminal domain (NTD), receptor binding domain (RBD), receptor binding motif (RBM), C-terminal domain (CTD), furin cleavage site (S1|S2), second proteolytic cleavage site (S2’), heptad repeats (HR), and transmembrane region (TMM). Domains were assigned according to the annotations from Cai, et al. (2020). **(B)** Phylogenetic profile of predicted epitopes; The taxa are ordered according to Fig. 1. Epi: Overlap with experimentally verified epitopes (Shrock, et al. 2020). Domain: The spike domain assignment for the LEs (c.f. (A)). Epitope losses are indicated by a green cell color. The occurrence of LE6 in the early branching sarbecovirus lineage (purple cell color) is due sto a convergent emergence of this epitope (cf. (C)). LE2 and LE6 are present only in 1/3 of the bat viruses subsumed under the respective node in the tree, which is indicated by a lighter cell color. See Fig. S7 for a version of this figure with taxon numbers provided. **(C)** Multiple sequence alignment for the nine predicted linear epitopes across representatives of the sarbecoviruses.

However, the spike protein is heavily glycosylated, and the glycans shield a considerable fraction of the protein from interactions with host molecules (Casalino, et al. 2020; Sikora, et al. 2020; Watanabe, et al. 2020; Wintjens, et al. 2020; Yang, et al. 2020). Since structural information was not considered in the prediction procedure, we integrated the position of the 9 epitopes with surface-accessibility information from a recent structural and physics-based modeling approach (Sikora, et al. 2020). This analysis suggests that LE3 is inaccessible in spike’s pre-fusion conformation (Fig. S8), while LEs 1, 4 and 7 are surface exposed but partially covered by glycans (Table S3). Complementary evidence for the accessibility of the linear epitopes comes from various experimental studies. All LEs, except for LE4, overlap with experimentally confirmed epitopes resulting from a study exploring the humoral response in Covid-19 patient sera (Shrock, et al. 2020)(Figs. 4B and S7). LE3 is a cross-reactive epitope that is identified by antibodies against the spikes of SARS-CoV-2 and of common cold viruses (HCoV) (Ng, et al. 2020). LE6 and LE8 overlap with the epitopes of two neutralizing human monoclonal antibodies against SARS-CoV-2, CB6 (Shi, et al. 2020) and 4A8 (Chi, et al. 2020), respectively, and LE7 is a “popular” epitope against which antibodies are detected in many Covid-19 patients (Shrock, et al. 2020). Taken together, there is sufficient evidence that the predicted LEs are exposed at least at some point during infection, and thus potentially participate in the interaction with the host.

Epitope scans in the spike orthologs revealed that the number of linear epitopes shared with the SARS-CoV-2 spike decreases with the evolutionary distance (Figs. 4 and S7). Secondary epitope losses are rare, with the exception of RmYN02, where a recombination event (Zhou, et al. 2020a) removed most of the ancestral epitopes. The LE analysis provides the first indication of a step-wise remodeling of the spike protein surface on the evolutionary lineage leading to SARS-CoV-2 (Fig. 5). The four epitopes located in the RBD (LE2,4-6), which establishes the connection to ACE2, are of particular interest. Three of them (LE2,5,6) overlap with predicted B-cell epitopes that are at most partly shielded by glycans (Wintjens, et al. 2020). LEs 2 and 4, which correspond to the N- and C-termini of the RBD are present in almost all sarbecoviruses. LE5, which directly flanks LE2, is confined to the SARS-CoV-2 clade. LEs 4 and 5 are absent in the spikes of the GX pangolin viruses due to substitutions of otherwise fully conserved amino acid (Fig. 4C).

**Figure 5.**
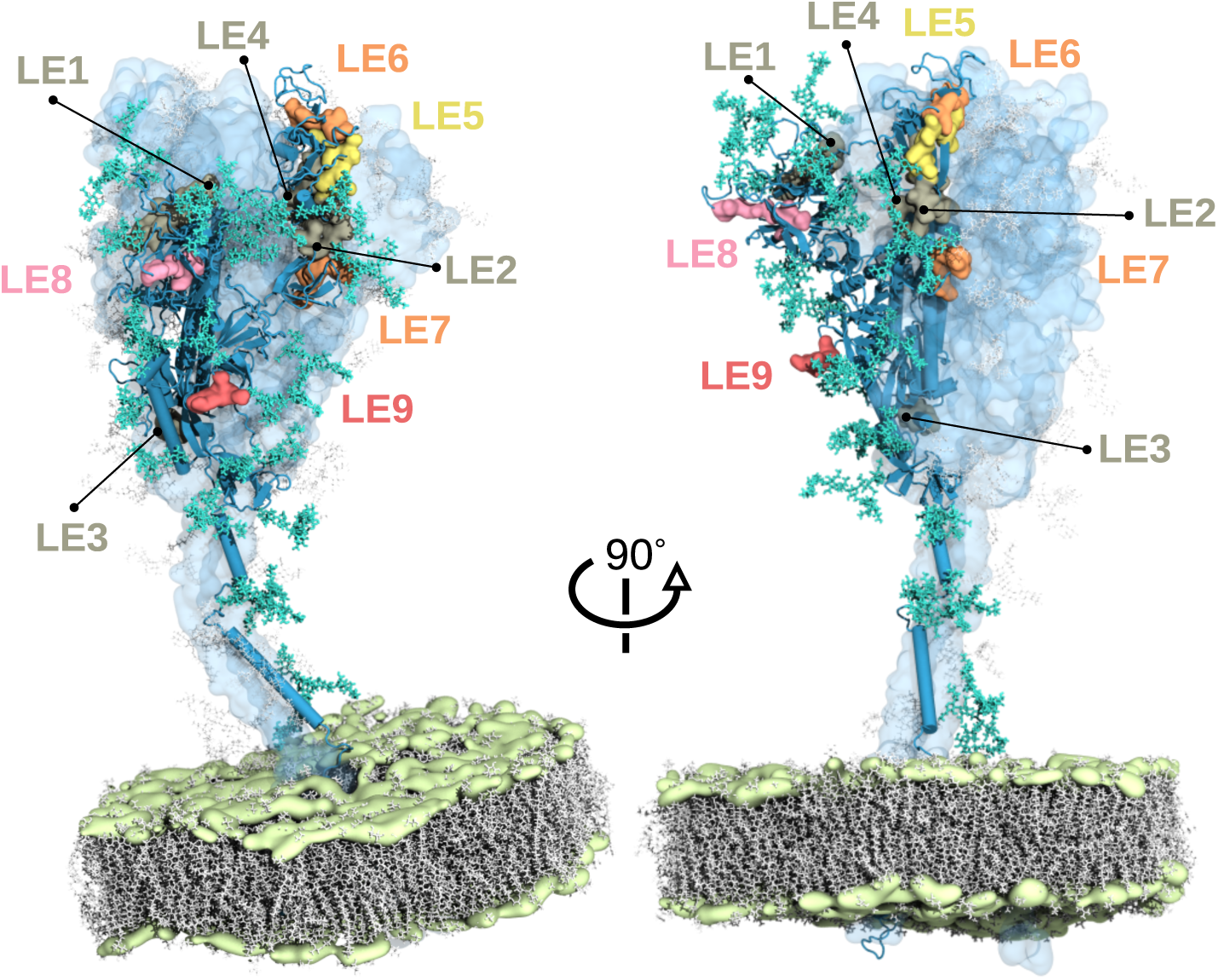
Predicted linear epitopes annotated on a full-length model of a spike glycoprotein of SARS-CoV-2. The structural model (Sikora, et al. 2020) represents the trimeric spike in a prefusion state embedded in a lipid bilayer (lipid tails marked gray and phosphate headgroups green). The epitopes in the surface representation are overlaid with the chain in the “up” conformation (blue cartoon). Realistic glycan coverage is shown as turquoise licorice. The remaining two chains in the “down” conformation are shown as light-blue transparent surfaces, with corresponding glycans transparent gray. The epitope color code follows Fig. 4B.

Together with the overall rarity of secondary epitope losses, it is tempting to speculate that these losses adapt the spike to the pangolin host. Notably, LE6 at the tip of the RBD (Figs. 5), which harbors four contacting residues to human ACE2 (Y449, Y453, L455, F456; (Lan, et al. 2020)) is present in RaTG13 and pangolin viruses. Thus, the virus was already preadapted to binding to human ACE2 long before its zoonotic transfer to humans.

Interestingly, LE6 is partly occluded when the RBD is down (Table S3; Fig. S9), which appears to be the preferential conformation of the spike in RatG13 and the pangolin viruses (Antoni, et al. 2021). It becomes fully exposed upon the conformational destabilization of the S trimer induced by the proteolytic cleavage (Wrobel, et al. 2020)(Fig. S9). Thus, LE9 – representing the furin cleavage site – allowed SARS-CoV-2 to carry into effect the binding capacity provided by an otherwise occluded RBM.

## Discussion

In the past two decades, two human pandemics, SARS and COVID-19, emerged from zoonotic transfers of sarbecoviruses. This sub-genus of the beta-coronaviruses infects a broad range of mammalian species. However, they are preferentially found in horseshoe bats (Zhou, et al. 2020b), and thus far only viruses isolated from bats span the full known phylogenetic diversity of the sarbecoviruses (our study, (Boni, et al. 2020; Zhou, et al. 2020a; Zhou, et al. 2020b)). Bats tolerate the infection with viruses that are lethal in other mammals (Schountz 2014). They found means to counteract the immune-modulatory effects of viral proteins, which otherwise result in a pro-inflammatory cytokine response that is associated with immunopathology, the main reason for severe and fatal outcomes of Covid-19 (Irving, et al. 2021; Perlman and Netland 2009) (for a review see (Banerjee, et al. 2020)). Consequently, bats presumably provide the main evolutionary playground for many pathogenic viruses, and in particular for sarbecoviruses (see Supplementary Discussion). Here we have investigated the innovations on the protein level that distinguish SARS-CoV-2 from other coronaviruses to shed light on the evolutionary trajectory transforming an animal virus into a human pathogen.

### Genetic innovation on the sub-domain level

The phylogenetic profiling of the SARS-CoV-2 protein set revealed that the sub-genus sarbecovirus is genetically distinct from all other coronaviruses. Differences in the feature architecture in particular of NSP1 and NSP3, which are present also outside sarbecovirus indicate a functional divergence of these proteins during coronavirus evolution. Within the sarbecoviruses, however, neither changes in gene set composition nor of feature domain architecture drive evolutionary diversification directing the attention to more subtle changes. Whole genome alignments of sarbecoviruses have meanwhile provided detailed information about sequence conservation/divergence patterns on a per-base resolution along a genome, and within coding regions in particular for the spike (Boni, et al. 2020; Li, et al. 2020b; Liu, et al. 2020). Yet, a substantial proportion of the spike is inaccessible either because the residues are buried or because they are shielded by glycans (Casalino, et al. 2020; Sikora, et al. 2020; Turonova, et al. 2020), such that most mutations might not be visible to the host. The functional relevance of sequence changes outside sites of known function is difficult to assess. Comparative studies on the protein structure level (e.g. Wrobel, et al. (2020)) integrated with molecular dynamics simulations (Sikora, et al. 2020) provide a more direct view on relevant changes and similarities. Yet, the computational demand of such approaches renders them inappropriate for comparative studies across many taxa. With the phylogenetic profiling of predicted linear epitopes we present an approach of intermediate resolution that can be used for rapidly scanning protein surfaces for lineage specific changes on a sub-domain level.

Molecular dynamics simulations considering the glycan shield (Sikora, et al. 2020) integrated with viral epitope profiling (Shrock, et al. 2020) indicate that except LE4 all predicted linear epitopes are visible to the host at some point during infection. LEs 1, 3, and 7 are B-cell immunodominant, although structural inspections suggest that they are in the case of LE3 fully, and of LEs 1 and 7 partly inaccessible (Table S3; Fig. S8). However, protein surface accessibility assessments are based on the spike in pre-fusion conformation (Sikora, et al. 2020). Some epitopes will be exposed by spike’s reorganization to the post-fusion conformation. Short-lived intermediate structures along this reorganization have thus far eluded both experimental and simulation approaches (Cai, et al. 2020; Wrobel, et al. 2020). The phylogenetic profiling of linear epitopes presented here therefore complements structure-based approaches. It has the potential to uncover “blind spots” of the structure-based accessibility analyses due to partial knowledge of the full conformational landscape.

### Evolution of the epitope landscape

The epitope landscape of the SARS-CoV-2 spike protein is the product of a gradual remodeling during sarbecovirus evolution (Figs. 4, 5). A comparison of SARS-CoV-1 with SARS-CoV-2 reveals that both pathogens display largely distinct sets of epitopes on their spike surfaces (Figs. S7 and S10). In fact, only few human antibodies against SARS-CoV-1 cross-react with SARS-CoV-2 (Pinto, et al. 2020), and they are largely confined to the S2 domain of the spike, which contains only one of our LEs (Ng, et al. 2020). The substantial overhaul of epitope landscapes on the two viral lineages underlines that SARS-CoV-2 is not just an evolutionarily refined version of SARS-CoV-1, consistent with their distinct placement in the virus phylogeny.

Our approach has one natural limitation, that is shared with experimental epitope profiling studies (Ng, et al. 2020; Shrock, et al. 2020). Structural epitopes that emerge from the spatial proximity of non-adjacent peptides in the amino-acid sequence in the folded protein are not considered. It is for this reason that we miss a shared epitope between SARS-CoV-1 and SARS-CoV-2 that is recognized by the same antibody (Pinto, et al. 2020). Although the restriction to linear epitopes comes at the cost of a reduced sensitivity, it renders our approach scalable and independent of the availability of accurate 3D-models of the proteins under study. It thus can be rapidly applied to any novel variant of SARS-CoV2, as well as on any newly emerging human pathogen.

### Epitope function balances immunodominance

The interaction of the virus with its host requires that the pathogen exposes parts of the spike epitopes—most prominently the RBD and the RBM therein—to establish contact with the ACE2 (Lan, et al. 2020; Walls, et al. 2020). The resulting epitopes should be highly immunogenic, and their evolutionary emergence and subsequent fate is determined by a trade-off between their function, and the counter-selective pressure imposed by the host immune system. For LE9, representing the furin cleavage site, and LE6, which harbors four contact residues to ACE2, the balance between risk and payoff of the epitope is clearly in favor of the latter. Both are critical for the infection process (Hoffmann, et al. 2020b; Lan, et al. 2020; McCallum, et al. 2020; Shang, et al. 2020a). It is still unclear why the other epitopes are maintained. In particular, the presence of LEs 3 and 8 should be strongly counter-selected. Both are detected by neutralizing human antibodies (Chi, et al. 2020; Ng, et al. 2020). LE8 is present also in RaTG13 and in viruses from the Guangdong pangolins, which suggests that selection could have acted long enough to eradicate the epitope. LE3, which is located in the S2 domain of the spike, is even more compelling. This epitope is present in all sarbecovirus spikes except that of RmYN02, and it shows cross-reactivity to antibodies formed against human CoV (Ng, et al. 2020). We see two possible explanations for the presence of these epitopes in the SARS-CoV-2 spike: Selection in bats and possible intermediate hosts was more permissive, for which indeed evidence exists (Banerjee, et al. 2020), and the evolutionary time under new and more rigorous selective regime in humans was too short to drive the loss of this epitope. Alternatively, LE8 and even more so the conserved LE3 may play essential roles in spike’s function that requires its presence despite opening a flank to the host’s immune system. It is tempting to speculate that the same is true for the other epitopes, in particular for LEs that are common to the spikes of all sarbecoviruses. Such evolutionarily stable epitopes could represent preferred targets for the design of therapeutic interventions against a broad spectrum of sarbecoviruses. In fact, targeting epitopes on the conserved stalk of influenza viruses is a promising strategy for a universal vaccine development (Nachbagauer, et al. 2021).

### Key innovations in the emergence of SARS-CoV-2

RaTG13 and the GD pangolin viruses share all but one linear epitope with SARS-CoV-2. Our finding integrates with the recent observation that the spike protein isolated from the GD pangolin virus binds human ACE2 with about 10-fold higher affinity than the pangolin ACE2 (Antoni, et al. 2021). Together, this suggests that the spike in the common ancestor of these viruses was largely pre-adapted to humans, and presumably only subtle adjustments were required to optimize for human infection.

The spike of the animal viruses appears to reside mainly in the closed conformation with all RBDs down (Antoni, et al. 2021; Zhang, et al. 2020b). This renders the RBM, which is represented in our analysis by LE6, largely inaccessible (cf. Table S3). Comparative cryo-EM analyses revealed that the un-cleaved spike trimer of SARS-CoV-2 is substantially more stable than its counterpart in RaTG13 (Wrobel, et al. 2020). This reinforces the blocking of the RBM (cf. Fig. S9) and reduces binding affinity of spike to ACE2. However, at the same time, it hides the contained epitopes from the host immune system (Shang, et al. 2020a).

With the furin cleavage site (LE9), SARS-CoV-2 has acquired a trait that has been repeatedly connected to pathogen virulence (Molloy, et al. 1999). Furin cleavage sites are sporadically occurring in CoV, e.g. in MERS (Gierer, et al. 2013; Millet and Whittaker 2014), and in the murine coronavirus (MHV) (de Haan, et al. 2004), both viruses with a high mortality. The functional consequences of furin cleavage are multi-facetted and thus far only partly understood. One of the most intriguing aspects is linked to the hiding of immunodominant epitopes in the RBM from the host immune system – a strategy known as “conformational masking” (Shang, et al. 2020a). This allows the virus to assume a ‘stealth mode’, where it has a reduced binding affinity to ACE2 but better evades immune response by the host. Furin cleavage has been associated with the exposure of RBMs and the epitopes therein as reinforcement of ACE2 binding and facilitator of cell entry (Benton, et al. 2020; Wrobel, et al. 2020), and the virus then rapidly enters the host cell. Of note, the role of the furin cleavage site in conformational masking seems to be different in the SARS-CoV-2 variant carrying the D614G substitution (Isabel, et al. 2020; Korber, et al. 2020). D614G results in a constitutively more open conformation of spike (Benton, et al. 2020; Gobeil, et al. 2021; Turonova, et al. 2020; Yurkovetskiy, et al. 2020). However, upon furin cleavage the D614G variant of spike has been reported to adopt preferentially a conformation where all three RBDs point down, with the consequence that their RBMs are masked and inaccessible for antibodies, but also for ACE2 (Gobeil, et al. 2021). It is tempting to speculate about the role of secreted furin (Vidricaire, et al. 1993) that may cleave spike even when it is not receptor bound, and of other proprotein convertases able to cleave at the multibasic furin site (Papa, et al. 2021). It will be interesting to see if and how this apparently reversed role of furin cleavage on conformational masking in the D614G variant affects the virus’ strategy to evade the host immune system.

The functional importance of the furin cleavage extends beyond conformational masking. Pre-cleaved spikes allow the virus to infect cell types with low expression of TMPRSS2 (Shang, et al. 2020a), and promote syncytia formation facilitating cell-cell transfer of the virus (Papa, et al. 2021). However, depending on the cell type the presence of the furin cleavage site can be also strongly counter-selected. For example, in veroE6 cells that are infected via the endosomal pathway and not via the cell membrane (Klein, et al. 2020), the furin cleavage site is rapidly lost or inactivated (Ogando, et al. 2020; Turonova, et al. 2020). This reveals that the virus adjusts its infection program, and also the role of the furin cleavage site, in a cell type dependent manner (Klein, et al. 2020).

The diverse functional roles of the furin cleavage site suggest that the selective regime underlying its evolution is complex, and processes to optimize its use are likely still in progress. This renders any mutations that further fine-tune the conformational dynamics of the spike particularly threatening, and this should be included into the monitoring and risk-assessment of novel SARS-CoV-2 variant. The recently detected variant B1.1.7, which is substantially more infective than other SARS-CoV-2 strains, may fall in this category. It harbors a mutation in LE9 covering the furin cleavage site (P681H) (Rambaut, et al. 2020).

## Conclusions

Phylogenetic profiling is a rapid and low-cost approach to embed a pathogen into its evolutionary background and to screen for genetic innovations that contributed to its emergence. Here we could show that the implicit scanning of protein surfaces, via the prediction of linear T-cell epitopes, provides the necessary resolution to capture how SARS-CoV-2 gradually evolved from its common ancestor shared with other sarbecoviruses. Our findings suggest that the epitope landscape of the S-protein, the primary interface of virus-host interaction, was largely molded in bats. Strategies to monitor and prevent the emergence of SARS-CoV-3 should therefore concentrate on species fulfilling at least two requirements: Their ACE2 receptor should be similar to that of humans, and their immune system is permissive enough to let the virus explore the epitope landscape of its surface proteins. In this context, the phylogenetic profiling of predicted linear epitopes can be routinely applied even when no protein structures are available, and it will be interesting to see how the extension of the catalogue beyond T-cell epitopes affects the resolution.

## Data Availability

All data underlying the phylogenetic profiles are available for download from https://applbio.biologie.uni-frankfurt.de/download/SARS-CoV-2/. The feature aware phylogenetic profiles can be interactively explored at https://applbio.biologie.uni-frankfurt.de/phyloprofilecorona/

## Acknowledgements

The authors would like to thank Ngoc Vinh Tran for support with the customization of PhyloProfile and for setting up the PhyloProfileCorona server, Dr. Florian E. Blanc, Sören von Bülow, and Michael Gecht for sharing scripts and structural models, Bardya Djahanschiri, Dr. Stefanie Ebersberger, and Dr. Ahmad Reza Mehdipour for helpful discussion. M.S. acknowledges support by the Austrian Science Fund FWF (Schrödinger Fellowship, 515 J4332-B28). M. S. and G. H. acknowledge support by the Max Planck Society. R. C. acknowledges support by the Frankfurt Institute for Advanced Studies. This research was supported by the research funding program Landes-Offensive zur Entwicklung Wissenschaftlich-ökonomischer Exzellenz (LOEWE) of the State of Hessen, Research Center for Translational Biodiversity Genomics (TBG) and the Centre for multi-scale modelling, analysis and simulation of biological processes (CMMS).

## Notes

https://applbio.biologie.uni-frankfurt.de/download/SARS-CoV-2/

### Competing Interest Statement

The authors have declared no competing interest.

### Summary of Updates

The discussion related to the gain of the furin claevage site has been adjusted to better reflect the diverse role of this site during SARS-CoV-2 infection.

