## Supplementary material for "The evolutionary making of SARS-CoV-2": Table S3 and Figs. S1-S10

### Supplementary information

#### Supplementary Discussion

##### The evolutionary origins of SARS-CoV-2

Bats represent important natural reservoirs of zoonotic viruses because they tolerate the infection with viruses that are lethal for other mammals, and infections typically take a very mild course with no apparent fitness decrease (Irving, et al. 2021; Schountz 2014). Key to this tolerance is a robust antiviral immune response in particular against RNA viruses. Bats found means to counteract the immune-modulatory effects of viral proteins, which otherwise result in a pro-inflammatory cytokine response that is associated with immunopathology, the main reason for severe and fatal outcomes of Covid-19 (Perlman and Netland 2009)(for a review see (Banerjee, et al. 2020)). As a consequence, bats presumably provide the main evolutionary playground of many pathogenic viruses, and in particular of sarbecoviruses. Our results indicate that both direct zoonotic transfer from horseshoe bats (Jaimes, et al. 2020) and a detour via pangolins as an intermediate host (Lam, et al. 2020) are conceivable. Thus far, RaTG13 is genetically the closest relative to SARS-CoV-2 (Jaimes, et al. 2020). However, the paraphyly of the GD and GX pangolin viruses, together with an epitope that is exclusively shared with the viruses of the GD lineage, reveal that pangolins have been infected twice independently by a virus related to SARS-CoV-2. The viral diversity within pangolins is far from being exhaustively explored (Liu, et al. 2020), and thus, a third and hitherto unsampled pangolin virus may have given rise to SARS-CoV-2. But are pangolins incubators for past and, more importantly future sarbevoviral human pathogens? Spikes of both pangolin viruses have a higher binding affinity to human ACE2 than to that of pangolin (Antoni, et al. 2021; Zhang, et al. 2020). A long-term propagation of the virus in pangolins is therefore likely to result in an optimization of the binding affinity by lineage-specific adaptations of the spike that render a subsequent transfer to humans unlikely. Two epitope losses in the RBD of the GX pangolin viruses due to the substitution of otherwise evolutionarily conserved residues may point in the direction of such an adaptation process. In addition, pangolins infected with SARS-CoV-2 severely suffer from respiratory disease (Liu, et al. 2020). Thus, even if SARS-CoV-2 was transferred from pangolins, they probably merely served as jumping hosts rather than contributing to shape this pathogen. This role would be similar to what has been concluded for civets in the context of SARS-CoV-1 (Li, et al. 2005b; Wu, et al. 2005). Besides pangolins, there is a broad range of other and domesticated mammalian species whose ACE2 receptors have higher binding affinities to the SARS-CoV-2 spike protein (Damas, et al. 2020). Cats, in particular, can be efficiently infected, show severe epithelial lesions, and can also transmit the virus via the airborne route (Shi, et al. 2020). The regular surveillance of these potential intermediate hosts, and monitoring and characterization of viral innovation in horseshoe bats are therefore relevant countermeasures to prevent the re-introduction of SARS-CoV-2 from a non-human reservoir, as it is repeatedly observed with MERS (Dudas, et al. 2018), and to reduce the risk of the future emergence of a SARS-CoV-3. The phylogenetic profiling of linear epitopes can be routinely applied in such surveys.

#### Supplementary Data

##### Data S1. SARS-CoV-2 phylogenetic profile

Orthologous groups generated in this study are provided for download from https://applbio.biologie.uni-frankfurt.de/download/SARS-CoV-2/Proteome. Provided are the files for an interactive inspection of the feature-aware phylogenetic profile of SARS-CoV-2 using PhyloProfile. Before running PhyloProfile, the custom Coronaviridae taxonomic information provided in the supplement within the data directory should replace the files that come with the default installation of PhyloProfile. To do so, please follow the guidelines below

1. Install Phyloprofile according to the installation requirements and instructions as described in the software's website (https://github.com/BIONF/PhyloProfile)
2. Start R and identify the location where the PhyloProfile package was installed by issuing the command *find.package("PhyloProfile")* in the R prompt. We refer to this path now as the $INSTALLATIONPATH
3. Place the taxonomy files provided with the SARS-CoV-2 data set https://applbio.biologie.uni-frankfurt.de/download/SARS-CoV-2/taxon_data.zip into the directory $INSTALLATIONPATH/PhyloProfile/data. This can be done with any file browser
4. Start up PhyloProfile from the R prompt. Once the PhyloProfile web interface appears, you can upload the provided files for visualization
    - Main input file: coronavirus.phyloprofile
    - Additional annotation input: coronavirus.domains
    - Order taxa by user defined tree: coronavirus.nwk
    - FASTA file optional input: coronavirus.fasta

   The phylogenetic profile is represented in the main plot. The feature architecture can be inspected for each data point in the detailed view, showing the feature architecture of the protein in SARS-CoV-2 on top, and its respective ortholog at the bottom.

##### Data S2. SARS-CoV-2 phylogenetic profile of polyprotein components.
Orthologous groups for the SARS-CoV-2 P1ab components are provided for download from https://applbio.biologie.uni-frankfurt.de/download/SARS-CoV-2/P1ab

#### Supplementary Tables

##### Table S1. Overview of the taxon set.
An overview of the sequence data and viral strains analyzed here is provided in the document TableS1_taxonset.xlsx

##### Table S2. SARS-CoV-2 protein feature prevalence in Polyprotein 1ab orthologs.
Table S2 is provided in the document TableS2_P1ab_featureprevalence.xlsx

Table S3. Epitope accessibility based on molecular dynamics simulations

| | | Ray (max) | Ray (mean) | Docking (max) | Docking (mean) |
|---|---|---|---|---|---|
| $1^{Up}$ | **LE1** | 0,33 | 0,13 | 0,13 | 0,05 |
| | **LE2** | 0,80 | 0,41 | 0,42 | 0,22 |
| | **LE3** | 0 | 0 | 0,02 | 0,01 |
| | **LE4** | 0,38 | 0,06 | 0,26 | 0,04 |
| | **LE5** | 0,46 | 0,20 | 0,23 | 0,11 |
| | **LE6** | 0,88 | 0,55 | 1,00 | 0,53 |
| | **LE7** | 0,17 | 0,04 | 0,03 | 0,02 |
| | **LE8** | 0,74 | 0,33 | 1,00 | 0,47 |
| | **LE9** | 1,00 | 0,74 | 1,00 | 1,00 |
| $0^{Up}$ | **LE1** | 0,32 | 0,11 | 0,13 | 0,05 |
| | **LE2** | 0,73 | 0,41 | 0,42 | 0,22 |
| | **LE3** | 0 | 0 | 0,02 | 0,01 |
| | **LE4** | 0,38 | 0,06 | 0,26 | 0,04 |
| | **LE5** | 0,36 | 0,16 | 0,14 | 0,07 |
| | **LE6** | 0,51 | 0,23 | 0,23 | 0,09 |
| | **LE7** | 0,17 | 0,04 | 0,02 | 0,01 |
| | **LE8** | 0,74 | 0,33 | 1,00 | 0,47 |
| | **LE9** | 1,00 | 0,73 | 1,00 | 1,00 |

$1^{up}$ and $0^{up}$ represent simulations with 1 or 0 RBDs pointing upwards. The accessibility of the epitopes was probed by illuminating the simulated spike with diffuse light (ray) and by rigid body docking of the antibody CR3022 Fab as described in Sikora, et al. (2020). Yellow and red cell color indicate only partly accessible ($0.01 < \text{Docking}_{mean} \leq 0.1$) and inaccessible linear epitopes, respectively. LE6 (shaded in grey) shows a pronounced change in accessibility in 0up vs. 1up conformation.

#### Supplementary Figures

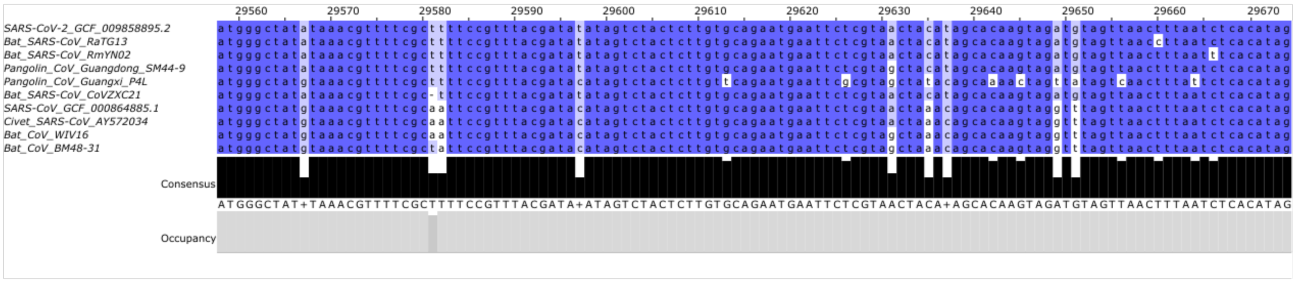

Figure S1. Multiple sequence alignment of the genomic region resembling SARS-CoV-2 ORF10 across representatives covering sarbecovirus diversity. The open reading frame is conserved in all analyzed taxa with the exception of BAT_SARS-CoV_CoVZXC21, where a deletion disrupts the reading frame.

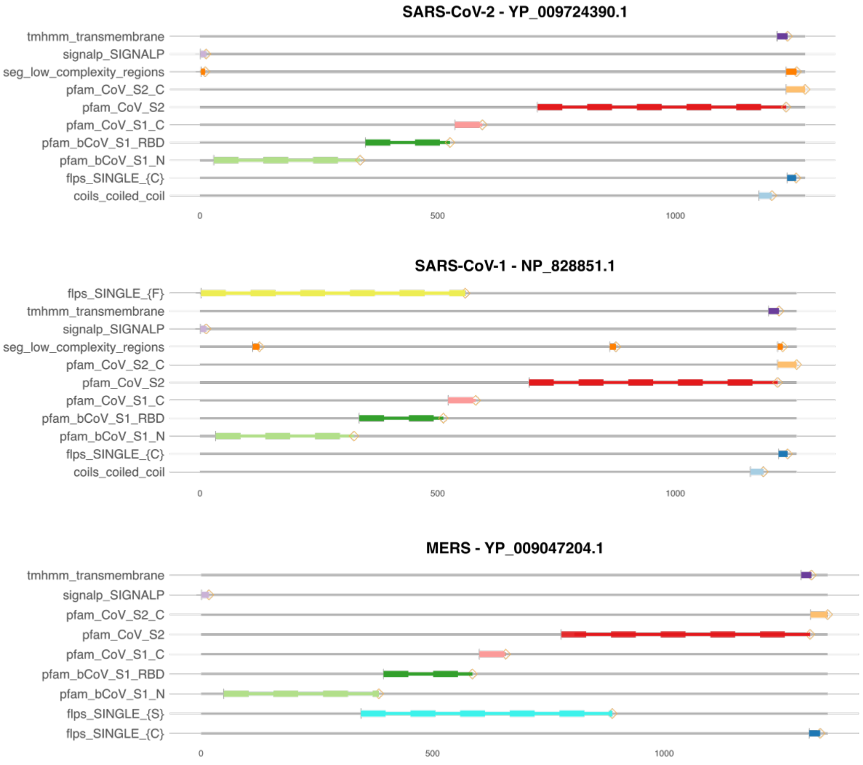

Figure S2. Feature architecture comparison for the spike glycoproteins of SARS-CoV-2, SARS-CoV-1, and MERS. Grey lines represent the amino acid sequences of the represented proteins, and colored bars indicate position and length of the protein parts occupied by the features indicated to the left of the bar. The x-axis gives the position in the amino acid sequence. The main layout of the architectures is similar for the three proteins, and differences are largely due to changes in amino acid composition. SARS-CoV-1 harbors a Phenylalanin (F)-rich N-terminal half that is not seen in SARS-CoV-2 or MERS, and the central region of the MERS spike is rich in Serine (S). Additionally, it lacks a coiled-coil region at the C-terminus, which is present in both SARS-CoV and SARS-CoV-2.

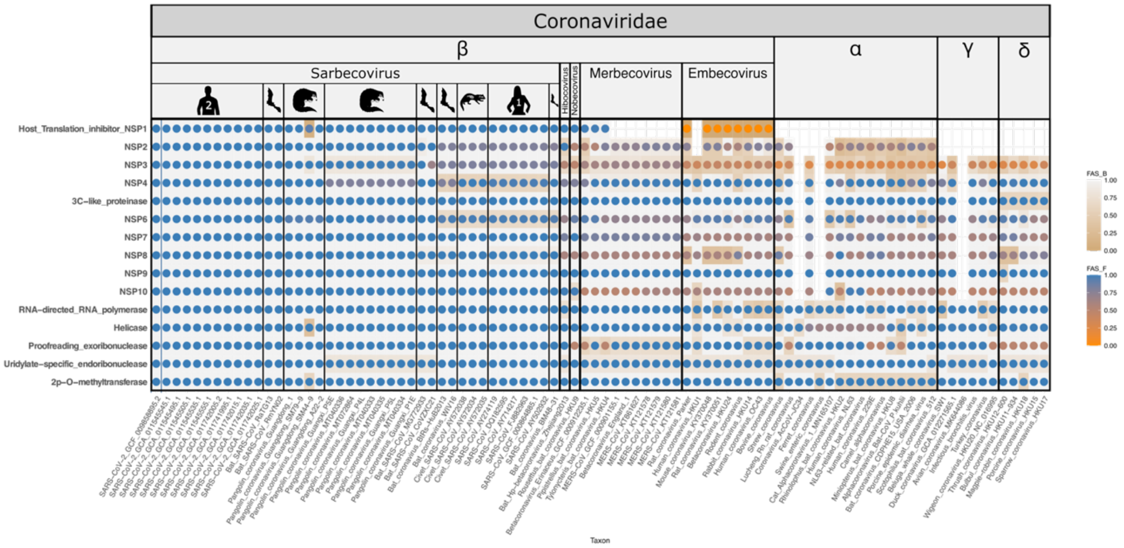

Figure S3. Phylogenetic profile of the 15 SARS-CoV-2 non-structural proteins (NPS) represented in the Polyprotein 1ab. Cleavage sites of the polyprotein components were assigned according to the entry P0DTD1 in UniProt for SARS-CoV-2, and retrieved for the taxonomic sampling using a pairwise alignment to the polyprotein 1ab orthologs. The presence of gaps in the profile indicate that the aligned regions were not identified as orthologs. The pronounced change in feature architecture in embecovirus NSP1 resembles the absence of the bCoV_NSP1 domain (PF11501), a domain that is otherwise present in all beta-CoV NSP1. Instead, embecovirus NSP1 harbor a Pfam domain of unknown function (DUF3477; PF11963). This change in architecture together with the pronounced sequence change of MERS NSP1 (see main text) underline the evolutionary plasticity of this protein both in sequence and function (Lokugamage, et al. 2015; Narayanan, et al. 2015)

```
SARS-CoV-2      ---------------MESLVPGFNEKTHVQLSLPVLQVRDVLVRG---FGDSVEEVLSEA
MERS            MSFVAGVTAQGARGTYRAALNSEKHQDHVSLTVPLCGSGNLVEKLSPWFMDGENAYEVVK
                                .: :  . :.: **.*::*:    ::: :    *  *. :

SARS-CoV-2      RQHLKDGTCGLVEVEKG-VLPQLEQP-YVFIKRSDARTAPHGHVMVELVAELEGIQYGR-
MERS            AMLLKKEPLLYVPIRLAGHTRHLPGPRVYLVERLIACENPFMVNQLAYSSSANGSLVGTT
                   **. .    * :. .    :*  *    :::*  *    *.    :   :. :*   *

SARS-CoV-2      -SGETLGVLVPHVGEIPVAYRKVLLRKNGNKGAGGHSYGADLKSFDLGDELGTDPYEDFQ
MERS            LQGKPIGMFFPYDIELVTGKQNILLRKYGR---GGYHYTP------FHYERDNTSCPEWM
                 .*:.:*::.*:   *:  .. :::**** *.    **: * .       :  * .. .  ::

SARS-CoV-2      ENWNTKHSSGVTRELMRELNGG
MERS            DDFEADPKGKYAQNLLKKLIGG
                :::::. ..  :::*:::* **
```

Figure S4. Pairwise sequence alignment of bCoV NSP1 reveals a conserved LLRKXG motif. SARS-CoV-2: Uniprot-Id P0DTC1; MERS: Uniprot-Id K9N638

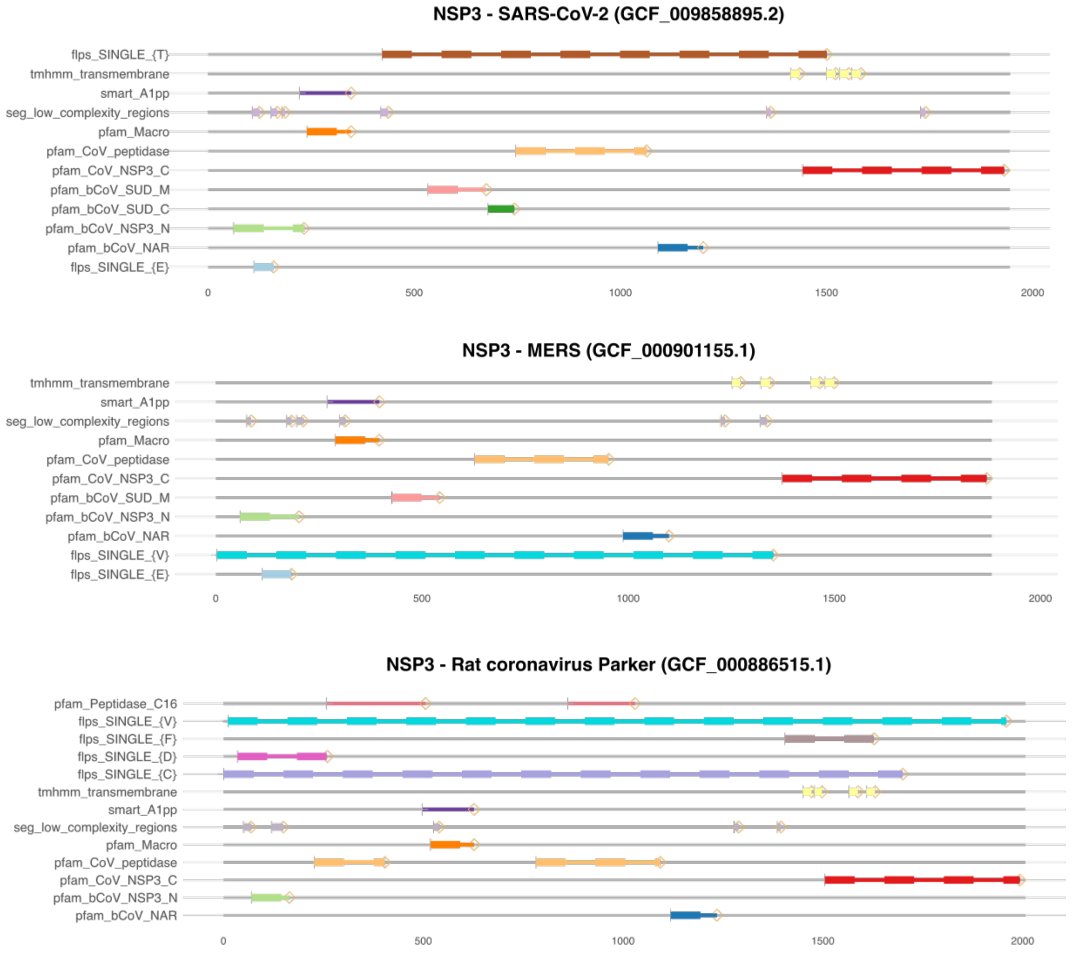

Figure S5. The evolutionary change of the NSP3 domain architecture during bCoV evolution. Grey lines represent the amino acid sequences of the represented proteins, and colored bars indicate position and length of the protein parts occupied by the features indicated to the left of the bar. The x-axis gives the position in the amino acid sequence. The three architectures represent NSP3 in sarbecovirus (top), merbecovirus (middle) and embecovirus (bottom). While the overall domain architecture is conserved throughout beta-CoV evolution, the N-terminal Pfam domain CoV_peptidase domain present in the embecovirus representative is absent in the other two sub-genera. In turn, the the Pfam domain bCoV_SUD_C is confined to sarbecoviruses.

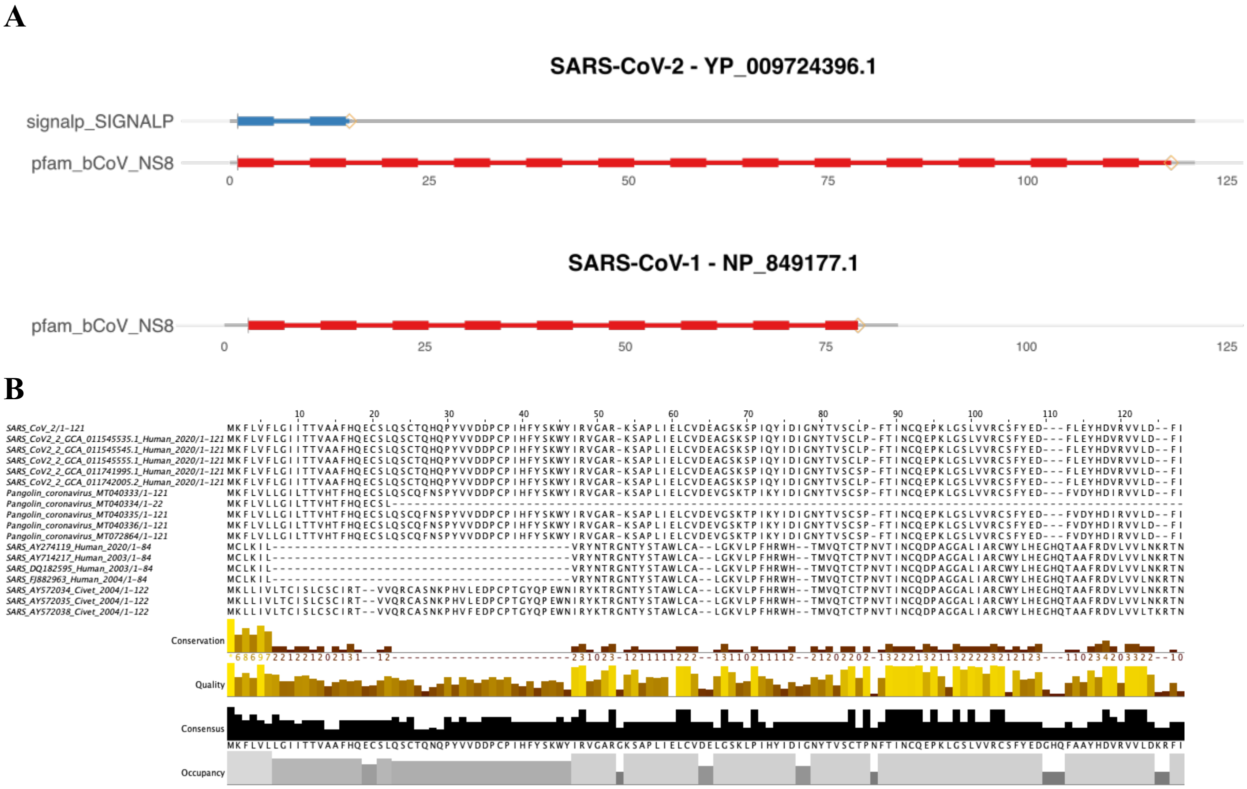

Figure S6. Loss of a signal peptide in SARS-CoV. A) Feature architecture comparison between ORF8 in SARS-CoV-2 and SARS-CoV, respectively. Grey lines represent the amino acid sequences of the represented proteins, and colored bars indicate position and length of the protein parts occupied by the features indicated to the left of the bar. The x-axis gives the position in the amino acid sequence. B) Multiple sequence alignment of ORF8 orthologs. The alignment reveals a large N-terminal deletion in the SARS-CoV-1 representatives resulting in the loss of the signal peptide.

| | Position | Epi | Domain | SARS-CoV-2 (175) | Bat_RaTG13 (1) | Bat_RmYN02 (1) | Pangolin_GD (4) | Pangolin_GX (9) | Bat_RecBP (2) | SARS-CoV-1 (6) | Civets (3) | SARS-CoV_bats Clade 1 (3) | SARS-CoV_bats Clade 2 (3) | Bat_root_SARS (1) | Hibecovirus (1) | Nobecovirus (2) | Outgroup (68) |
|---|---|---|---|---|---|---|---|---|---|---|---|---|---|---|---|---|---|
| **LE1** | 112-132 | 115 - 123 | NTD | 1 | 1 | 1 | 1 | 1 | 1 | 1 | 1 | 1 | 1 | 1 | 0 | 0 | (0,04) |
| **LE2** | 329-338 | 329 - 338 | RBD | 1 | 1 | 1 | 1 | 1 | 1 | 1 | 1 | 0,33 | 0 | 1 | 0 | 0 | (0,01) |
| **LE3** | 896-904 | 899-907 | S2 | 1 | 1 | 0 | 1 | 1 | 1 | 1 | 1 | 1 | 1 | 1 | 0 | 0 | 0 |
| **LE4** | 510-528 | - | RBD | 1 | 1 | 1 | 1 | 0 | 1 | 1 | 1 | 1 | 1 | 1 | 0 | 0 | 0 |
| **LE5** | 337-353 | 348-358 | RBD | 1 | 1 | 0 | 1 | 0 | 1 | 0 | 0 | 0 | 0 | 0 | 0 | 0 | (0,01) |
| **LE6** | 448-457 | 452-463 | RBM | 1 | 1 | 0 | 1 | 1 | 0 | 0 | 0 | 0 | (0,33) | (1) | 0 | 0 | (0,03) |
| **LE7** | 560-569 | 559-569 | CTD1 | 1 | 1 | 0 | 1 | 1 | 0 | 0 | 0 | 0 | 0 | 0 | 0 | 0 | 0 |
| **LE8** | 258-267 | 260 - 273 | NTD | 1 | 1 | 0 | 1 | 0 | 0 | 0 | 0 | 0 | 0 | 0 | 0 | 0 | (0,03) |
| **LE9** | 680-688 | 680-688 | S1/S2 | 1 | 0 | 0 | 0 | 0 | 0 | 0 | 0 | 0 | 0 | 0 | 0 | 0 | 0 |

Figure S7. The phylogenetic profile of predicted linear epitopes in the spike glycoprotein of SARS-CoV-2. We searched the spike proteins of 13 taxa for the presence of the SARS-CoV-2 epitopes. The number of proteins analyzed for each taxon are provided in parenthesis next to the taxon name. Values in the matrix represent the fraction of spike proteins harboring the epitope. The taxa are arranged according to an increasing evolutionary distance to SARS-CoV-2. Background color indicates the evolutionary history of the epitope: blue - conserved epitopes; green - secondary loss on the respective lineage; purple - convergent emergence. The accessibility was computed according to Sikora, et al. (2020). Parentheses indicate partially hidden LEs; Epi: Position of experimentally verified B cell epitopes based on a triple-Alanine scanning mutagenesis (Shrock, et al. 2020). The assignment of epitopes to Domains follows Cai, et al. (2020).

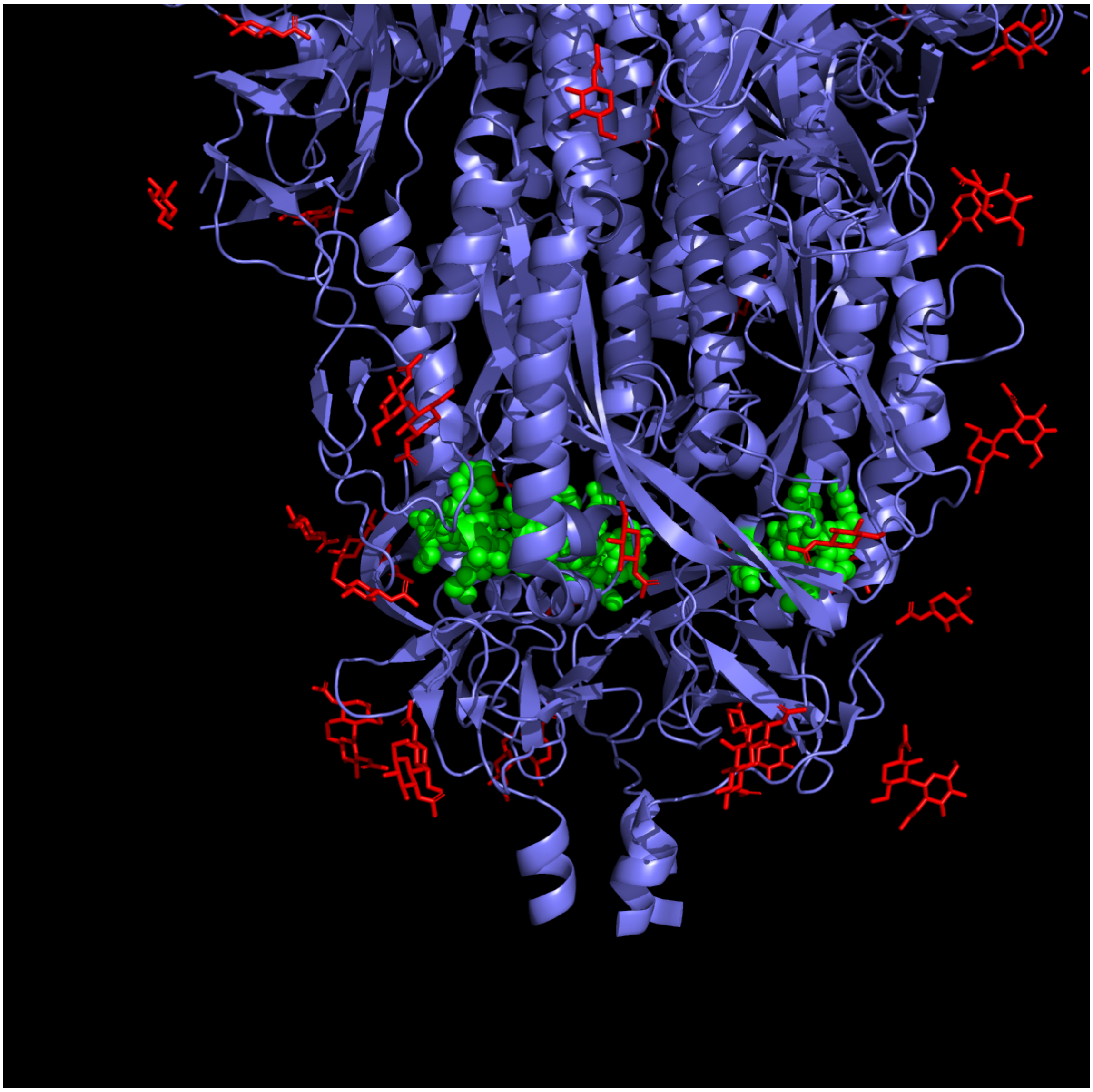

Figure S8 Masking of LE3. LE3 (green spheres) is nearly fully protected in the interior of a pre-fusion spike structure (PDBid 7kj3) and further masked by glycans (red).

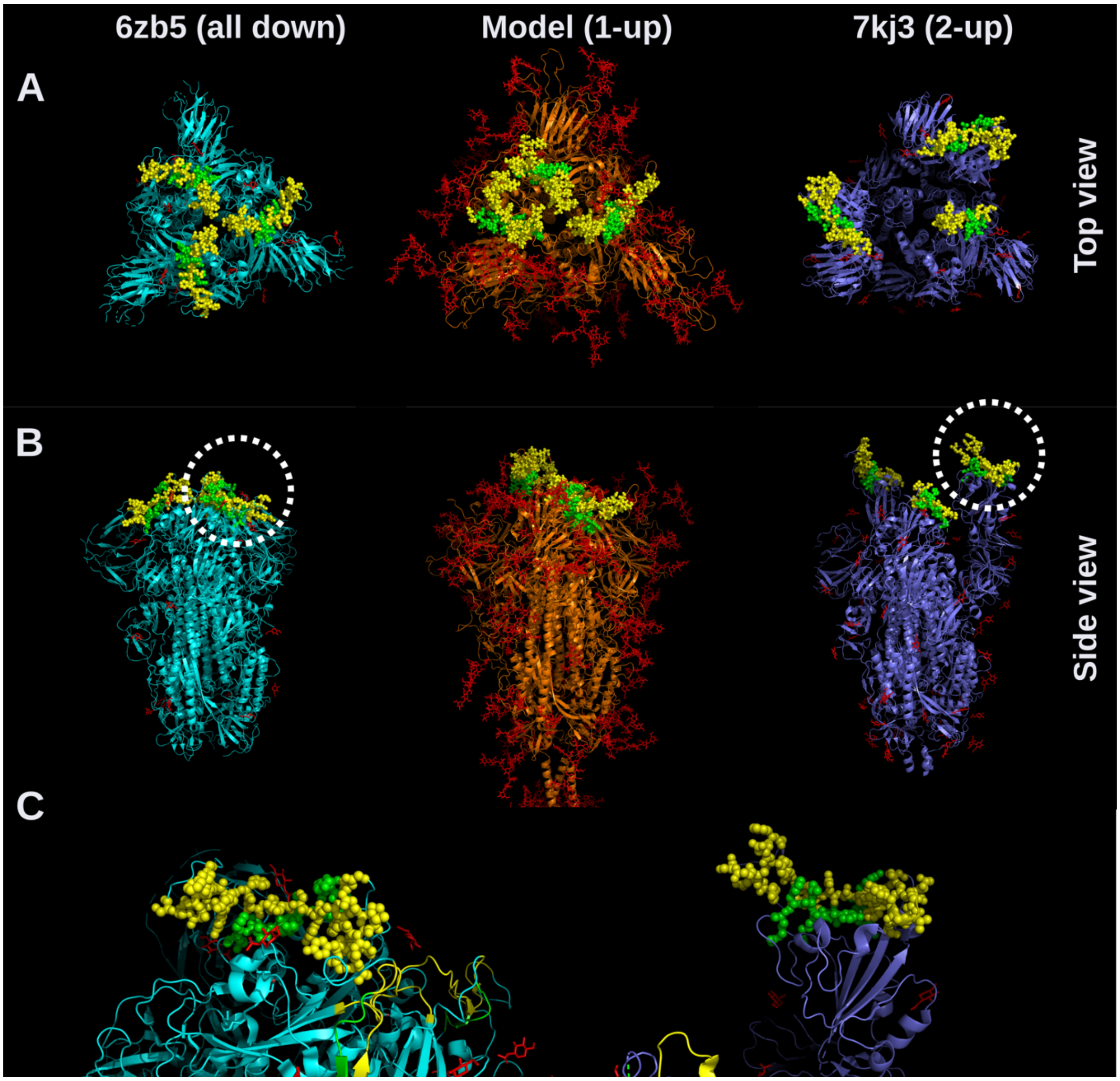

Figure S9. Location of LE6 in the spike. The panels show top (A) and side (B) views of different configurations of the spike protein: all-down (cyoEM structure, PDBid 6zb5); 1-up (full-length model; (Sikora, et al. 2020)); 2-up (cryoEM structure, PDBid 7kj3). LE6 (marked as green spheres) is nearly fully embedded within the ACE2 binding interface of the spike (yellow spheres)(Mehdipour and Hummer 2020). The exposure in the down conformation is reduced by the presence of glycans (red sticks), and it is enhanced by the opening of the RBDs. For cryoEM structures only the initial N-acetylglucosamine residues of a subset of glycans are visible. C) close up of an RBD in down and in up position, respectively. Access to the ACE2-binding interface and to LE6 therein is hindered in the closed conformation and is exposed in the up conformation.

| LE | Position | Domain | SARS-CoV-1 (6) | Civets (3) | SARS-CoV_bats Clade1 (3) | SARS-CoV_bats Clade2 (3) | SARS-CoV-2 (175) | Bat_RaTG13 (1) | Bat_RmYN02 (1) | Pangolin_GD (4) | Pangolin_GX (9) | Bat_RecBP (2) | Bat_root_SARS (1) | Hibecovirus (1) | Nobecovirus (2) | Outgroup (68) |
|---|---|---|---|---|---|---|---|---|---|---|---|---|---|---|---|---|
| **1** | 109 - 129 | NTD | 1 | 1 | 1 | 1 | 1 | 1 | 1 | 1 | 1 | 1 | 1 | 0 | 0 | (0,04) |
| **2** | 496 - 514 | RBD | 1 | 1 | 1 | 1 | 1 | 1 | 1 | 1 | 0 | 1 | 1 | 0 | 0 | (0,02) |
| **3** | 878 - 886 | S2 | 1 | 1 | 1 | 1 | 1 | 1 | 0 | 1 | 1 | 1 | 1 | 0 | 0 | 0 |
| **4** | 316 - 325 | RBD | 1 | 1 | 0,33 | 0 | 1 | 1 | 1 | 1 | 1 | 1 | 1 | 0 | 0 | (0,02) |
| **5** | 208 - 225 | NTD | 1 | 1 | 0,66 | 0 | 0 | 0 | 0 | 0 | 0 | 0 | (1) | 0 | 0 | 0 |
| **6** | 851 - 860 | S2 | 1 | 1 | 1 | 1 | 0 | 0 | 0 | 0 | 0 | 0 | 0 | 0 | 0 | 0 |
| **7** | 33 - 41 | NTD | 1 | 1 | 1 | 0,33 | 0 | 0 | 0 | 0 | 0 | 0 | 0 | 0 | 0 | (0,02) |
| **8** | 94 - 101 | NTD | 1 | 1 | 0,33 | 0 | 0 | 0 | (1) | 0 | 0 | 0 | 0 | 0 | 0 | (0,03) |
| **9** | 1 - 15 | SignalPep | 1 | 1 | 0,33 | 0 | 0 | 0 | 0 | 0 | 0 | (1) | 0 | 0 | 0 | (0,03) |
| **10** | 42 - 151 | NTD | 1 | 1 | 0 | 0 | 0 | 0 | 0 | 0 | 0 | 0 | 0 | 0 | 0 | 0 |
| **11** | 430 - 447 | RBM | 1 | 1 | 0 | 0 | 0 | 0 | 0 | 0 | 0 | 0 | 0 | 0 | 0 | 0 |
| **12** | 225 - 243 | NTD | 1 | 1 | 0 | 0 | 0 | 0 | 0 | 0 | 0 | 0 | 0 | 0 | 0 | 0 |
| **13** | 351 - 366 | RBD | 1 | 0 | 0 | 0 | 0 | 0 | 0 | 0 | 0 | 0 | 0 | 0 | 0 | (0,06) |

Figure S10. The phylogenetic profile of predicted linear epitopes in the spike glycoprotein of SARS-CoV-1. We searched the spike proteins of 13 taxa for the presence of the SARS-CoV-2 epitopes. The number of proteins analyzed for each taxon are provided in parenthesis next to the taxon name. Values in the matrix represent the fraction of spike proteins harboring the epitope. The taxa are arranged according to an increasing evolutionary distance to SARS-CoV-1. Background color indicates the evolutionary history of the epitope: blue - conserved epitopes; green - secondary loss; purple - convergent emergence. The assignment of epitopes to Domains follows Li, et al. (2005a).
